# Robust phylodynamic analysis of genetic sequencing data from structured populations

**DOI:** 10.1101/2022.06.16.496390

**Authors:** Jérémie Scire, Joëlle Barido-Sottani, Denise Kühnert, Timothy G. Vaughan, Tanja Stadler

**Affiliations:** Department for Biosystems Science and Engineering, ETH Zürich, Basel, 4058, Switzerland; Swiss Institute of Bioinformatics, 1015 Lausanne, Switzerland; Transmission, Infection, Diversification and Evolution Group, Max Planck Institute for the Science of Human History, 07745 Jena, Germany; Institut de Biologie de l’ENS, École normale supérieure, CNRS, INSERM, Université PSD, 75005 Paris, France

**Keywords:** phylogenetics, Bayesian inference, phylodynamics, population structure

## Abstract

The multi-type birth-death model with sampling is a phylodynamic model which enables quantification of past population dynamics in structured populations, based on phylogenetic trees. The BEAST 2 package *bdmm* implements an algorithm for numerically computing the probability density of a phylogenetic tree given the population dynamic parameters under this model. In the initial release of *bdmm*, analyses were limited computationally to trees consisting of up to approximately 250 genetic samples. We implemented important algorithmic changes to *bdmm* which dramatically increase the number of genetic samples that can be analyzed, and improve the numerical robustness and efficiency of the calculations. Including more samples leads to improved precision of parameter estimates, particularly for structured models with a high number of inferred parameters. Furthermore, we report on several model extensions to *bdmm*, inspired by properties common to empirical datasets. We apply this improved algorithm to two partly overlapping datasets of Influenza A virus HA sequences sampled around the world, one with 500 samples, the other with only 175, for comparison. We report and compare the global migration patterns and seasonal dynamics inferred from each dataset. In that way, we show what information is gained by analyzing the bigger dataset which became possible with the presented algorithmic changes to *bdmm*. In summary, *bdmm* allows for robust, faster and more general phylodynamic inference of larger datasets.

## 1. Introduction

Genetic sequencing data taken from a measurably evolving population contain fingerprints of past population dynamics [1]. In particular, the phylogeny spanning the sampled genetic data contains information about the mixing pattern of different populations and thus contains information beyond what is encoded in classic occurrence data, see e.g. Hey and Machado [2], Stadler and Bonhoeffer [3]. Phylodynamic methods [4,5] aim at quantifying past population dynamic parameters, such as migration rates, from genetic sequencing data. Such tools have been widely used to study the spread of infectious diseases in structured populations, see e.g. Dudas *et al*. [6], Faria *et al*. [7] as examples for analyses of recent epidemic outbreaks. The Bayesian phylodynamic inference framework BEAST2 [16] is one of the software frameworks within which such analyses can be carried out. With BEAST2, tree topologies, parameters from phylodynamic, molecular clock and substitution models can be jointly inferred via Markov-Chain Monte-Carlo sampling. Both the host population and the pathogen population may be structured, e.g. the host population may be geographically structured, and the pathogen population may consist of a drug-sensitive and a drug-resistant subpopulation. Understanding how these subpopulations interact with one another, whether they are separated by geographic distance, lifestyles of the hosts, or other barriers, is a key determinant in understanding how an epidemic spreads. In macroevolution, different species may also be structured into different “subpopulations”, e.g. due to their geographic distribution or to trait variations, see e.g. Hodges [8]. Phylodynamic tools aim at quantifying the rates at which species migrate or traits are gained or lost, and the rates of speciation and extinction within the “subpopulations”, see e.g. Goldberg *et al*. [9], Mayrose *et al*. [10], Goldberg *et al*. [11].

The phylodynamic analysis of structured populations can be performed using two classes of models, namely coalescent-based and birth-death-based approaches. Both have their unique advantages and disadvantages [12,13]. Here, we report on major improvements to a multi-type birth-death-based approach.

A multi-type birth-death model is a linear birth-death model accounting for structured populations. Under this model, the probability density of a phylogenetic tree can be calculated by numerically integrating a system of differential equations. The use of this model within a phylodynamic setting and the associated computational approach were initially proposed for analyzing species phylogenies [14] and later for analyzing pathogen phylogenies [12,15]. The BEAST2 package *bdmm* generalizes the assumptions of these two initial approaches [17]. It further allows for co-inferring phylogenetic trees together with the model parameters and thus takes phylogenetic uncertainty explicitly into account. Datasets containing more than 250 genetic sequences could not be analyzed using the original *bdmm* package [17] due to numerical instability. This limitation was a strong impediment to obtaining reliable results, particularly for analysis of structured populations, as quantifying parameters which characterize each subpopulation requires a significant amount of samples from each of them. The instability was due to numerical underflow in the probability density calculations, meaning that probability values extremely close to zero could not be accurately calculated and stored. We have solved the numerical instability issue of *bdmm*, thereby lifting the hard limit on the number of samples that can be analyzed. In addition, the practical usefulness of the *bdmm* package was previously restricted by the amount of computation time required to carry out analyses. We report here on significant improvements in computation efficiency. As a result, *bdmm* can now handle datasets containing several hundred genetic samples. Finally, we made the multi-type birth-death model more general in several ways: homochronous sampling events can now occur at multiple times (not only the present), individuals are no longer necessarily removed upon sampling, and the migration rate specification has been made more flexible by allowing for piecewise-constant changes through time.

Overall, these model generalizations and implementation improvements enable more reliable and ambitious empirical data analyses. Below, we use the new release of *bdmm* to quantify Influenza A virus spread around the globe as an example application, and compare the results obtained with those from the reduced dataset analyzed in Kühnert *et al*. [17].

## 2. Methods

### Description of the extended multi-type birth-death model

First, we formally define the multi-type birth-death model on *d* types [17] including the generalizations introduced in this work. The process starts at time 0 with one individual, this is also called the origin of the process and the origin of the resulting tree. This individual is of type *i* ∈ {1 … *d*}, with probability *h*_*i*_ (where 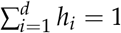). The process ends after *T* time units (at present). The time interval (0, *T*) is partitioned into *n* intervals through 0 < *t*_1_ < … < *t*_*n*−1_ < *T*, and we define *t*_0_ := 0 and *t*_*n*_ := *T*. Each individual at time *t, t*_*k*−1_ ≤ *t* < *t*_*k*_, *k* ∈ {1 … *n*} of type *i* ∈ {1 … *d*}, gives rise to an additional individual of type *j* ∈ {1 … *d*}, with birth rate *λ*_*ij,k*_, migrates to type *j* with rate *m*_*ij,k*_ (with *m*_*ii,k*_ = 0), dies with rate *μ*_*i,k*_, and is sampled with rate *ψ*_*i,k*_. At time *t*_*k*_, each individual of type *i* is sampled with probability *ρ*_*i,k*_. Upon sampling (either with rate *ψ*_*i,k*_ or probability *ρ*_*i,k*_), the individual is removed from the infectious pool with probability *r*_*i,k*_. We summarize all birth-rates *λ*_*ij,k*_ in ***λ***, migration rates *m*_*ij,k*_ in ***m***, death rates *μ*_*i,k*_ in ***μ***, sampling rates *ψ*_*i,k*_ in ***ψ***, sampling probabilities *ρ*_*i,k*_ in ***ρ*** and removal probabilities *r*_*i,k*_ in ***r***, *i, j* ∈ {1, …, *d*}, *k* ∈ {1, …, *n*}. The model described in Kühnert *et al*. [17] is a special case of the above, assuming that migration rates are constant through time (i.e. do not depend on *k*), removal probabilities are constant through time and across types (i.e. do not depend on *k* and *i*), and that *ρ*_*i,k*_ = 0 for *k* < *n* and *i* ∈ {1 … *d*}.

This process gives rise to complete trees on sampled and non-sampled individuals with types being assigned to all branches at all times (Figure 1, left). Following each branching event, one offspring is assigned to be the “left” offspring, and one the “right” offspring, each assignment has probability 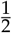. In the figure, we draw the branch with assignment “left” on the left and the branch with assignment “right” on the right. Such trees are called oriented trees, and considering oriented trees will facilitate calculations of probability densities of trees. Pruning all lineages without sampled descendants leads to the *sampled phylogeny* (Figure 1, middle and right). The orientation of a branch in the sampled phylogeny is the orientation of the corresponding branch descending the common branching event in the complete tree. When the sampled phylogeny is annotated with the types along each branch, we refer to it as a *branch-typed tree* (Figure 1, middle). On the other hand, if we discard these annotations but keep the types of the sampled individuals, we call the resulting object a *sample-typed (or tip-typed) tree* (Figure 1, right).

**Figure 1.**
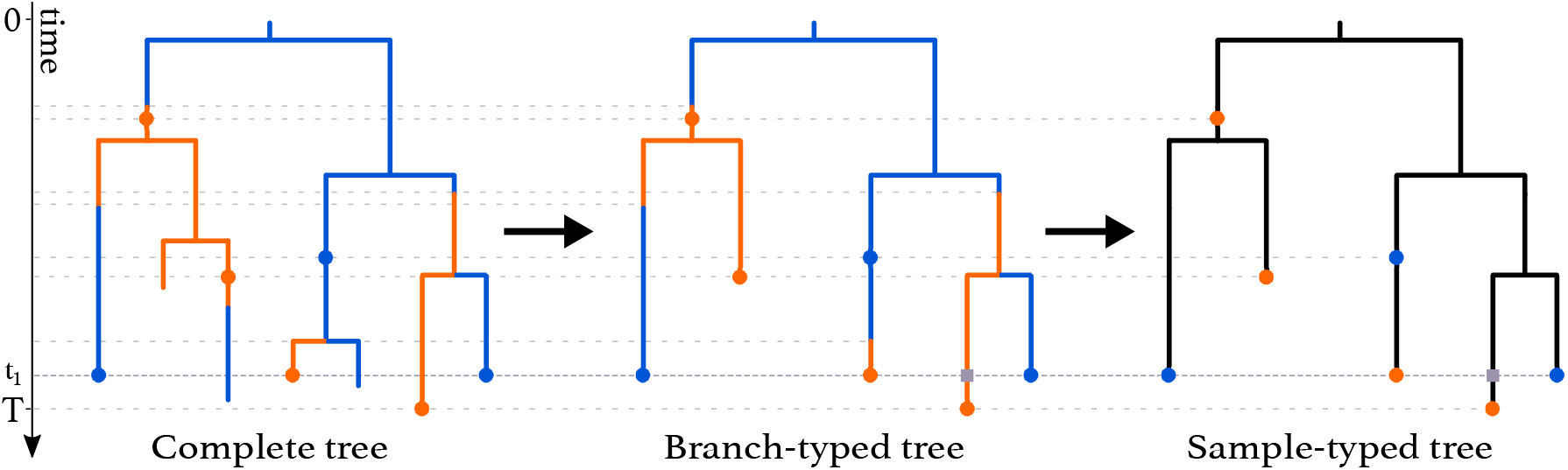
Complete tree (**left**) and sampled trees (**middle** and **right**) obtained from a multi-type birth-death process with two types. The orange and blue dots on the trees represent sampled individuals and are colored according to the type these individuals belong to. A *ρ*-sampling event happens at time *t*_1_. The grey squares represent degree-2 nodes added to branches crossing this event. *ρ*-sampling also happens at present (time *T*). As seen in the complete tree, the three individuals who were first sampled were not removed from the population upon sampling whereas the three individuals sampled at time *t*_1_ were.

Here, we give an overview of the computation of the probability density of the sampled tree (i.e. the sample-typed or branch-typed tree) given the multi-type birth–death parameters ***λ, m, μ, ψ, ρ, r, h***, *T*. This probability density is obtained by integrating probability densities *g* from the leaf nodes (or “tips”), backwards along all edges (or “branches”), to the origin of the tree. Our notation here is based on previous work [17,18], and the probabilities *p*_*i,k*_(*t*) and 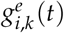 relate to *E* and *D* in Maddison *et al*. [14], Stadler and Bonhoeffer [15], respectively.

Every branching event in the sampled tree gives rise to a node with degree 3 (i.e. 3 branches are attached). Every sampling event gives rise to a node of degree 2 (called “sampled ancestor”) or 1 (called “tip”, as defined above). A sampling event at time *t* = *t*_*k*_, *k* ∈ {1, …, *n*}, is referred to as a *ρ*-sampled node. All other nodes corresponding to samples are referred to as *ψ*-sampled nodes.

Further, degree-2 nodes are put at time *t*_*k*_ on all lineages crossing time *t*_*k*_, *k* = 1, …, *n* − 1 as shown at time *t*_1_ in Figure 1. In a branch-typed tree, a node of degree 2 also occurs on a lineage at a time point when a type-change occurs. Such type changes may be the result of either migrations or birth events in which one of the descendant subtrees is unsampled (Figure 1, middle).

We highlight that in *bdmm*, we assume that the most recent sampling event happens at time *T*. This is equivalent to assuming that the sampling effort was terminated directly after the last sample was collected, and overcomes the necessity for users to specify the time between the last sample and the termination of the sampling effort at time *T*.

The derivation of the probability density of a sampled tree under the extended multi-type birth-death model is developed in Appendix A. This probability density, also called “phylodynamic likelihood”, can be used to estimate multi-type birth-death parameters ***λ, μ, ψ, m***, *T* when used in a Bayesian phylodynamic framework, such as BEAST 2 Bouckaert *et al*. [16]. Note that, unlike other parameters of this model, ***h*** is typically not estimated via MCMC sampling. *h*_*i*_ values can be set according to different rationales: the root type can be fixed to a particular type *k* (*h*_*k*_ = 1, *h*_*i*_ = 0 for *i* ≠ *k*), or all types can be equally likely 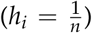 or they can be set to the equilibrium distribution (derived by Stadler and Bonhoeffer [15]) given the process was already in equilibrium at the time of origin.

### Implementation improvements

The computation of probability densities of sampled trees under the multi-type birth-death model require numerically solving Ordinary Differential Equations (ODEs) along each tree branch. We significantly improved the robustness of the original *bdmm* implementation, which suffered from instabilities caused by underflow of these numerical calculations. Compared to the original implementation, we prevent underflow by implementing an extended precision floating point representation (EPFP) for storing intermediary calculation results. Additional to this improvement in stability, we improved the efficiency of the probability density calculations, by 1) using an adaptive-step-size integrator for numerical integration, 2) performing preliminary calculations and storing the results for use during the main calculation step and 3) distributing calculations among threads running in parallel. Details can be found in Appendix B.

The latest release with our updates, *bdmm* 1.0, is freely available as an open access package of BEAST 2. The source code can be accessed at https://github.com/denisekuehnert/bdmm.

## 3. Results

### Evaluation of numerical improvements

We compared the robustness and efficiency of the improved *bdmm* package against its original version. We considered varying tree sizes, between 10 and 1000 samples. For each tree size, we simulated 50 branch-typed and 50 sample-typed trees under the multi-type birth-death model using randomly-drawn parameter values from the distributions shown in Table A1. The distributions from which parameters are drawn were selected to reflect a wide-range of scenarios. For each simulated tree, we measured the time taken to perform the calculation of the probability density given the parameter values under which the tree was simulated, using the updated and the original *bdmm* implementation. We report the wall-clock time taken to perform this calculation 5000 times (Figure 2). All computations are performed on a MacBook Pro with a dual-core 2.3 GHz Intel Core i5 processor. The new implementation of *bdmm* is on average 9 times faster than the original (Figure 2a). The robustness of the updated implementation is demonstrated by reporting how often the implementations return −∞ for the probability density in log space. We call these calculations “failures”, the most likely cause of error being underflow. Our new implementation shows no calculation failure for trees containing a thousand samples, while in the original implementation calculations often fail for trees with more than 250 samples (Figure 2b).

**Figure 2.**
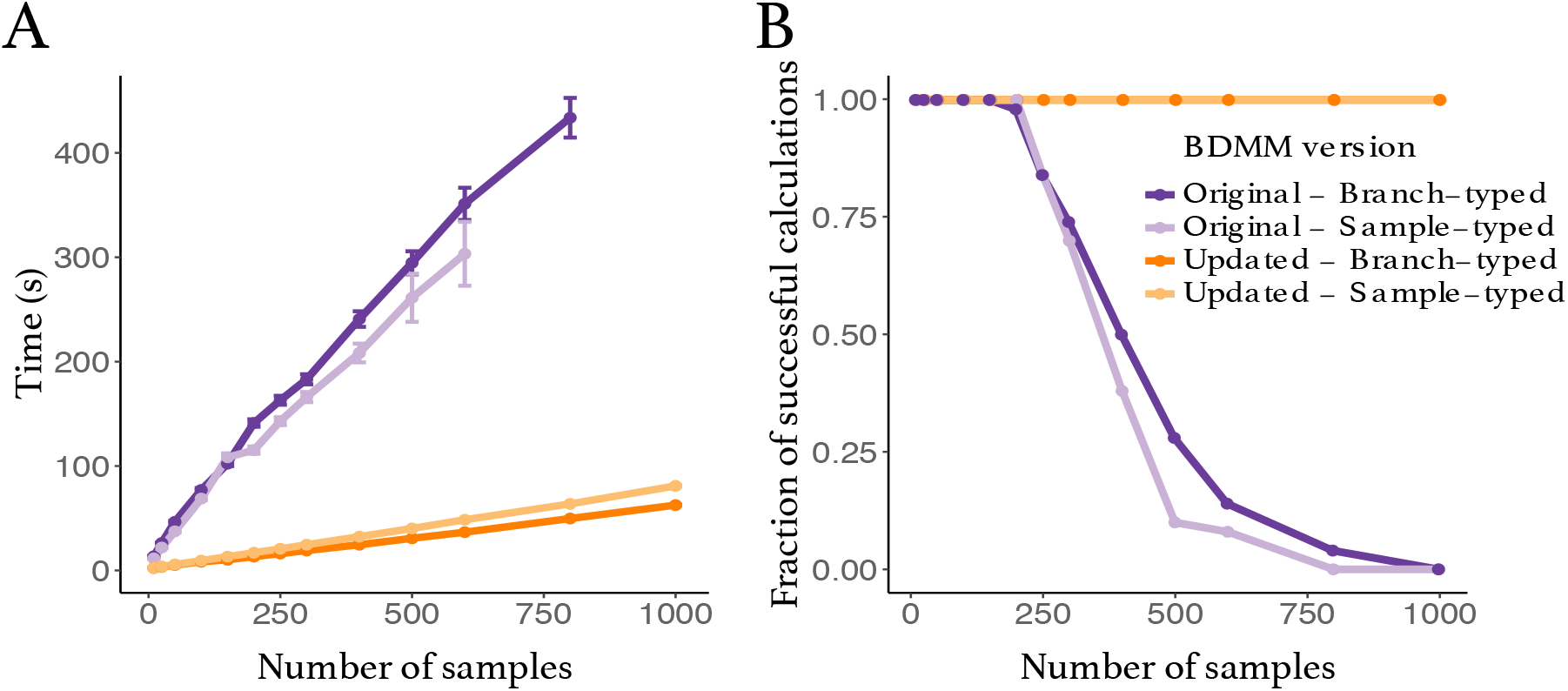
Comparison between the original and updated implementation of the multi-type birth-death model. (**a**) Speed comparison. Only successful calculations were taken into account i.e. calculations where the log-probability density was different from −∞. (**b**) Success in calculating probability density values plotted against tree size. The values presented in this panel correspond to the same set of calculations as the one in panel **a**.

### Validation against original implementation

To ensure that no errors were introduced into the updated *bdmm*, we validated the improved implementation against the original *bdmm* version by comparing results of likelihood computation on a handful of randomly-simulated trees. We used simulated trees of 10 or 100 tips, well below the limit of reliability of the original *bdmm* version (approximately 250 tips). Details of this procedure can be found in Appendix C. Fig. 3 shows one of such simulated trees along with tree likelihood values (or probability density of the sampled tree given the multi-type birth–death parameters) computed with each *bdmm* version. Likelihood computations results are identical for all trees and parameters tested between both implementations (difference in log-likelihoods < 1 × 10^−6^). Fig. A3 shows that the same results were obtained with other trees, or when varying other parameters. Therefore, we conclude that the results of the full validation, along with error and bias assessment, performed by Kühnert *et al*. [17] on the original *bdmm* version holds true for the improved *bdmm* implementation we present in this article.

**Figure 3.**
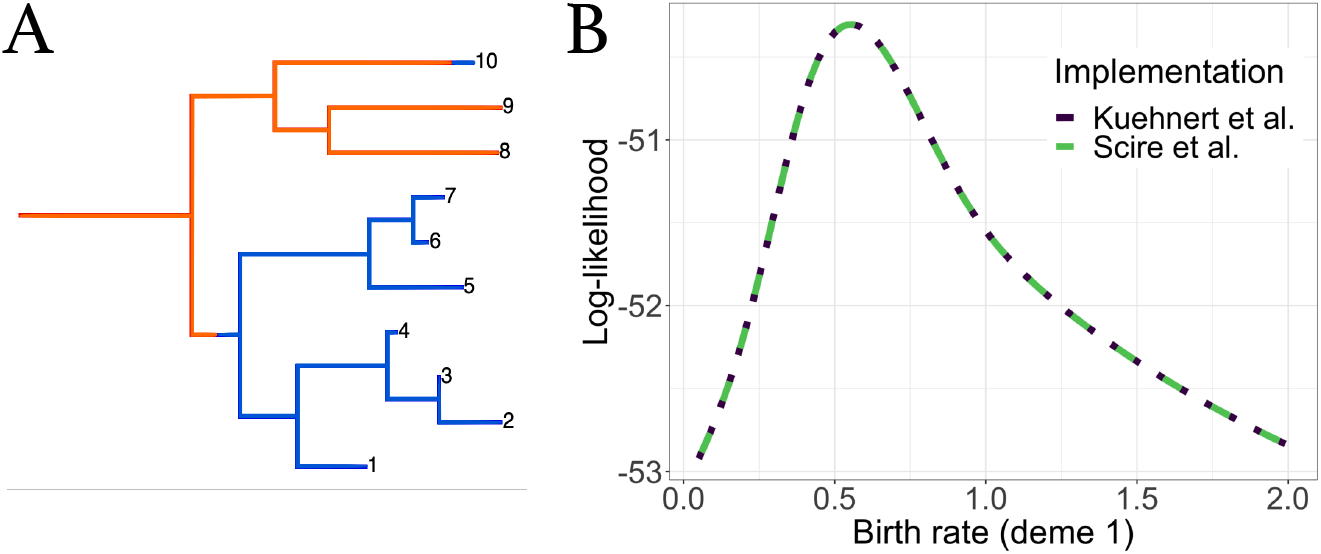
Comparison of computation results between the original *bdmm* and improved *bdmm* versions. **(a)** Randomly-simulated tree with 10 tips and 2 demes, used for comparison. (**b**) Log-likelihood values obtained with each *bdmm* version as a function of *λ*_1_ (birth rate of orange deme).

### Influenza A virus (H3N2) analysis

As an example of a biological question which can be investigated with *bdmm*, we analyzed 500 H3N2 influenza virus HA sequences sampled around the globe from 2000 to 2006 and aim to recover the seasonal dynamics of the global epidemics. The dataset is a random subset of the data analyzed by Vaughan *et al*. [19], taken from three different regions around the globe: New York (North, *n* = 167), New Zealand (South, *n* = 215) and Hong Kong (Tropics, *n* = 118). The dataset of 980 samples assembled by Vaughan *et al*. [19] was built with the aim of gathering samples from three locations with relatively-similar population sizes, each representative of the northern, southern or equatorial regions.

As a comparison, we performed an identical analysis with the H3N2 influenza dataset of 175 sequences sampled between 2003 and 2006 used in [17]. This dataset of 175 sequences was also a subset of the data by Vaughan *et al*. [19], and it also groups samples from 3 locations denoted as North (for northern hemisphere), South (for southern hemisphere) and Tropic (for tropical regions). Note that the latter dataset comes from more geographically-spread samples and thus we do not expect results from both analysis to be perfectly comparable. As we deal with pathogen sequence data, we adopt the epidemiological parametrization of the multi-type birth-death model as detailed in Kühnert *et al*. [17]. The epidemiological parametrization substitutes birth, death and sampling rates with effective reproduction numbers within types, rate at which hosts become noninfectious and sampling proportions. To study the seasonal dynamics of the global epidemic, we allow the effective reproduction number *R*_*e*_ to vary through time. To do so, we subdivide time into six-month intervals (starting April 1st and October 1st) and we constrain effective reproduction number values corresponding to the same season across different years to be equal for each particular location, assuming that the *R*_*e*_ values are the same in summer seasons and the same in winter seasons. Testing the validity of this hypothesis by estimating *R*_*e*_ values which vary in each 6-month period is not done as we expect little information from the data for the additional parameters. For the same reason, the migration rate is not varied through time. Further details on the data analysis configuration can be found in Appendix D.

The analysis of the larger dataset (500 samples) allows for the reconstruction of the phylogenetic tree encompassing a longer time period, and therefore gives a more long-term and detailed view of the evolution of the global epidemic (see Figure 4 for the Maximum-Clade Credibility trees).

**Figure 4.**
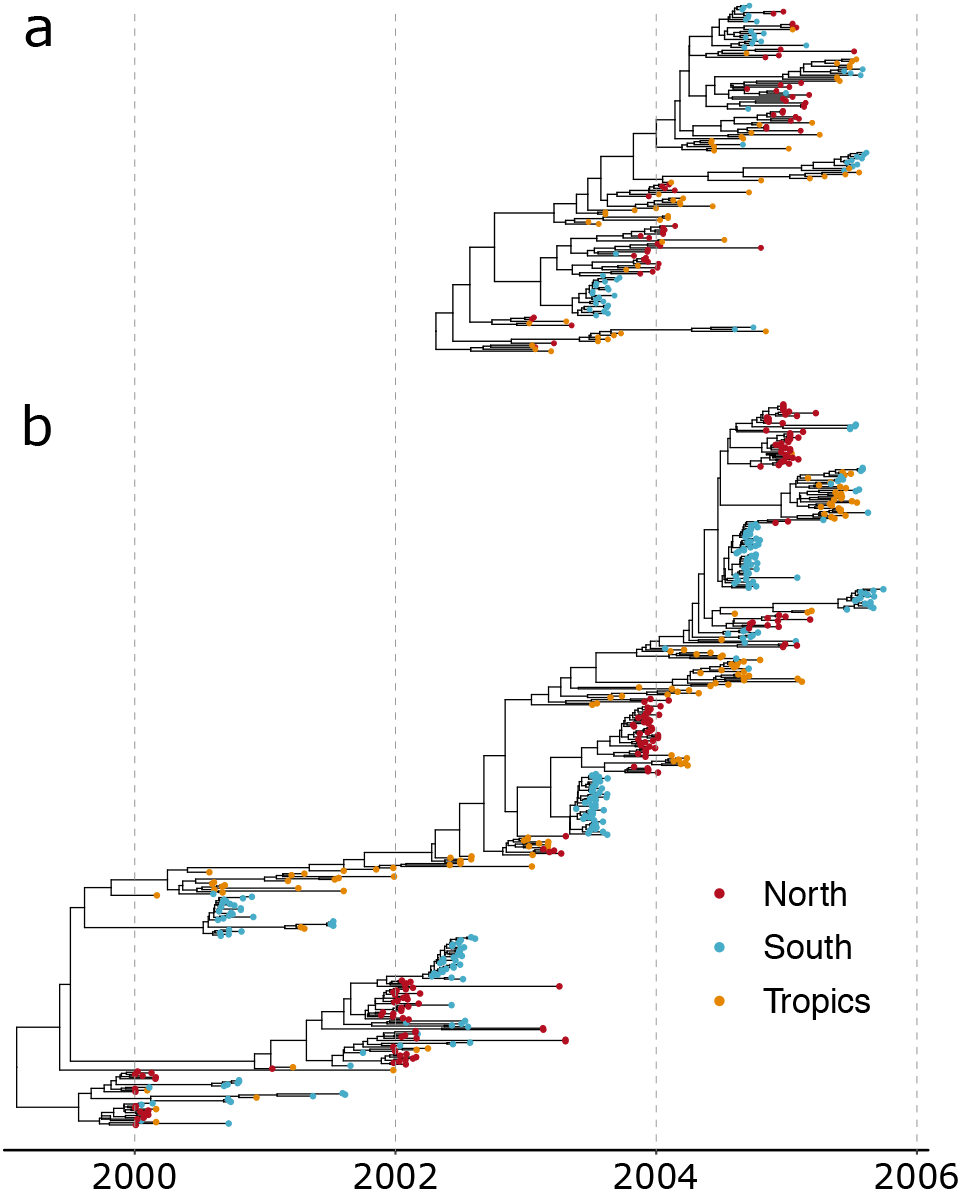
Maximum-Clade Credibility (MCC) trees for analyses with (**a**) 175 samples and (**b**) 500 samples.

As can be expected for the tropical location, in both analyses, effective reproduction numbers for H3N2 influenza A are inferred to be close to one year-round (Figure 5a). Conversely, strong seasonal variations can be observed in Northern and Southern hemisphere locations. There, the effective reproduction number is close to one in winter, while it is much lower in summer. Inferences from the small and large datasets are mostly in agreement. A subtle difference appears: in the larger dataset, the effective reproduction number in winter seasons and in the tropical location are closer to one, with less variation across estimates. This seems to indicate that the variations between estimates observed with the smaller dataset including samples from 2003 to 2006 (for instance *R*_*e*_ in winter in the North compared to *R*_*e*_ in winter in the South) are due to stochastic fluctations which are averaged out when considering a longer period of transmission dynamics in the larger dataset covering the years 2000 to 2006.

**Figure 5.**
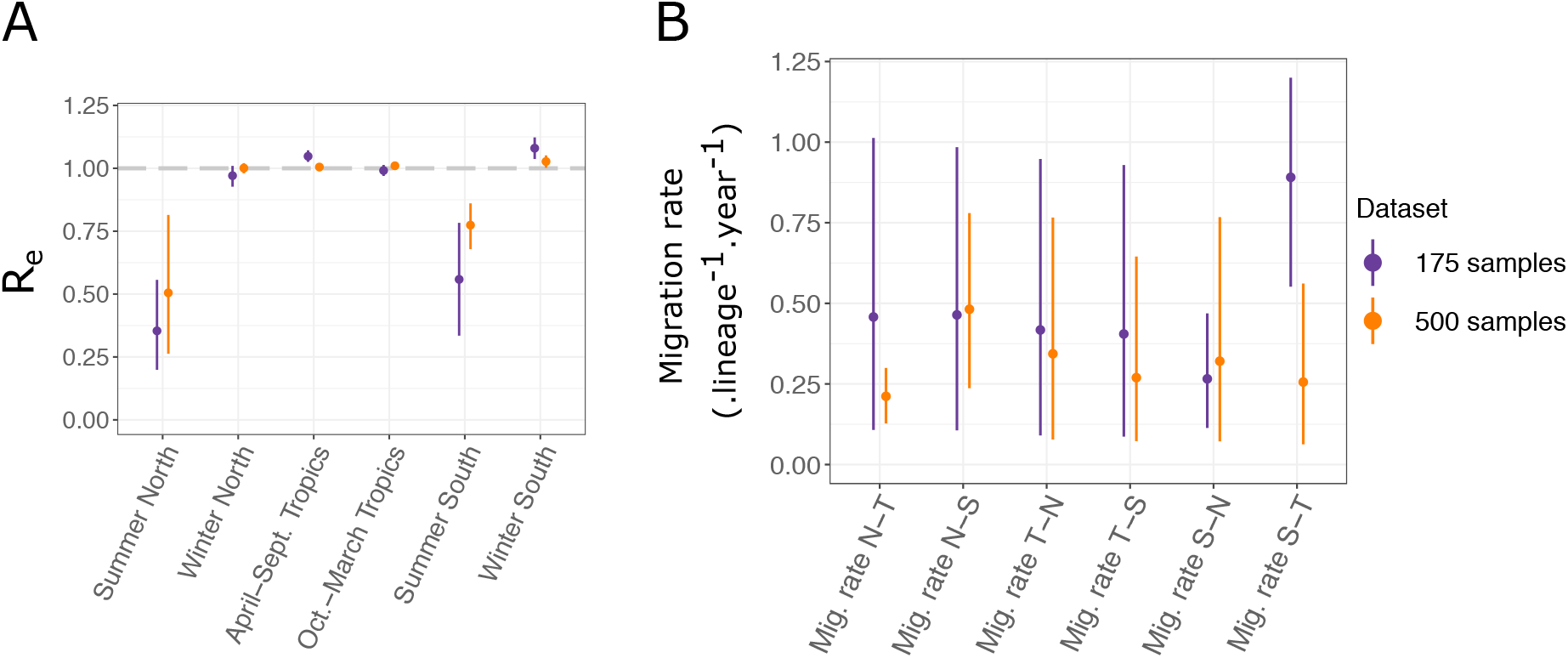
(**a**) Seasonal effective reproduction numbers for each sample location, for both datasets. (**b**) Migration rates inferred for each dataset. N, T and S refer respectively to North, South and Tropics. For instance, “*Mig. rate N-T*” represents the migration rate from the Northern location to the Tropical one.

Precise inference of migration rates is more difficult, as is reflected by the significant uncertainty we obtain on the estimates (Figure 5b). Still, we observe in general that the uncertainty is reduced for the inference performed with the larger dataset, as expected. A significant difference between the migration rates inferred from the Southern to Tropical locations arises between the two analysis. With the larger dataset, the estimated rate is much lower than with the smaller one, and more in range with the other migration rate estimates. Detailed results of all the parameter estimates for both analyses are available in Table A4. Most notably, estimates of root locations for both datasets are very similar. In both cases, the tropical location is the most likely location for the root; however, neither of the two other locations can be entirely excluded.

### *Properties of* bdmm

#### Identifiability of parameters

In birth-death models with sampling through time or at present, only two of the three parameters: birth rate, death rate, sampling rate/sampling probability can be jointly estimated [20,21]. Thus, independent prior data needs to be employed to quantify all three parameters. In recent work, the question of identifiabiliity in time-dependent birth-death-sampling models has been thoroughly investigated [21,22]. The issue of identifiability in state-dependent birth-death-sampling models remains, to our knowledge, largely unanswered. The interactions between migration rates, rates of birth-among-demes and other multi-type birth–deathparameters is not well-known. It is likely that different parameter combinations of the multi-type birth–deathmodel can yield the same likelihood value. Informative prior information on some of the birth-death parameters mitigates parameter non-identifiability issues.

#### Computational costs

Despite the implementation improvements presented in this manuscript, phylodynamics analyses performed in *bdmm* are still limited in practice in the number of genetic sequences they can handle. This limitation, unlike the previously-existing limitation caused by underflow, is not a hard boundary, but rather a soft boundary imposed by practical constraints of computational analyses. Limits in the complexity of analyses that can be carried out with the improved version of *bdmm* derive from the time required to carry out computations and from the complexity of the probability space that must be explored. For instance, each MCMC chain for the 500-sample Influenza A analysis required about 15 days to compute. Keeping the same analysis setup and increasing the number of genetic samples will have a linear effect on the time required to compute the phylodynamic likelihood with *bdmm*. With our updated *bdmm* implementation, the core bottleneck is the complexity of exploring tree space which increases exponentially with additional samples. Due to this complexity, only trees up to around 1000 samples can be estimated with BEAST2 successfully.

## 4. Discussion

The multi-type birth-death model with its updated implementation in the *bdmm* package for BEAST 2 provides a flexible method for taking into account the effect of population structure when performing phylodynamic genetic sequence analysis. Compared to the original implementation, we now prevent underflow of numerical calculations and speed up calculations by almost an order of magnitude. The size limit of around 250 samples for datasets which could be handled by *bdmm* is thus lifted and significantly larger datasets can now be analyzed. Now, the bottleneck is the search of tree space with MCMC rather than *bdmm*. We demonstrate this improvement by analysing two datasets of Influenza A virus H3N2 genetic data from around the globe. One dataset has 500 samples and could not have been analyzed with the original version of *bdmm*, the other one contains 175 samples and is the original example dataset analyzed in [17]. Overall, we observe, as expected, that analysing a dataset with more samples gives a more long-term picture of the global transmission patterns and reduces the general uncertainty on parameter estimates.

With the addition of so-called *ρ*-sampling events in the past, intense sampling efforts limited to short periods of time (leading to many samples being taken nearly simultaneously) can be easily modelled as instantaneous sampling events across the entire population (or sub-population), rather than as non-instantaneous sampling over small sampling intervals. This simplifies the modelling of intense pathogen sequencing efforts in very short time windows. By allowing the removal probability *r* (the probability for an individual to be removed from the infectious population upon sampling) to be type-dependent and to vary across time intervals, as well as allowing migration rates between types to vary across time intervals, we further increase the generality and flexibility of the multi-type birth-death model. An example *bdmm* analysis with *ρ*-sampling event in the past was added to the software package to guide users who may want to set up such an analysis with their own data.

We focused on an epidemiological application of *bdmm*, where we co-infer the phylogenetic trees to take into account the phylogenetic uncertainty. However, the *bdmm* modelling assumptions are equally applicable to the analysis of macroevolutionary data, in which context *bdmm* allows for the joint inference of trees with fossil samples under structured models [23]. When using a multi-type birth-death model in the macroevolutionary framework, *ρ*-sampling can be used to model fossil samples originating from the same rock layer. In the context of the exploration of trait-dependent speciation, structured birth-death models such as the binary-state speciation and extinction model (BiSSE) [24,25] have been shown to possibly produce spurious associations between character state and speciation rate when applied to empirical phylogenies [26]. When used in this fashion, users of *bdmm* should assess the propensity for Type I errors with their dataset through neutral trait simulations, as suggested by Rabosky and Goldberg [26].

In summary, the new release of *bdmm* overcomes several constraints when analysing sequencing data in BEAST. As it stands, the main constraint now is given by the efficiency of the BEAST2 MCMC tree space sampler rather than *bdmm* itself. We expect the new release of *bdmm* to become a standard tool for phylodynamic analysis of sequencing data and phylogenetic trees from structured populations.

## Supplementary Materials

The XML files for replication of the BEAST2 analyses of the two datasets of respectively 175 and 500 samples of H3N2 influenza virus HA sequences are available as supplementary files. To run these XML files using BEAST2, the *bdmm* and *feast* packages must be installed.

## Author Contributions

Conceptualization, J.S., J.B.-S., D.K., T.G.V. and T.S.; software, J.S., D.K. and T.G.V.; writing–original draft preparation, J.S., T.G.V and T.S.; writing–review and editing, J.S., J.B.-S., D.K., T.G.V. and T.S.; supervision, T.S.

## Acknowledgments

We thank Nicola Müller, David Rasmussen, Jū lija Pečerska for very valuable discussions and Fábio Kuriki Mendes for helpful comments on the manuscript.

## Appendix A. Derivation of the probability density of a sampled tree

### Appendix A.1. Probability of having no sampled descendants

In order to compute the probability density of a sampled tree, we need to calculate the probability of an individual with type *i* ∈ {1 … *d*} at time *t*_*k*−1_ ≤ *t* ≤ *t*_*k*_ to have no sampled descendant, which we denote by *p*_*i,k*_(*t*). In the interval *t*_*k*−1_ ≤ *t* < *t*_*k*_, this probability satisfies the ordinary differential equation (ODE)

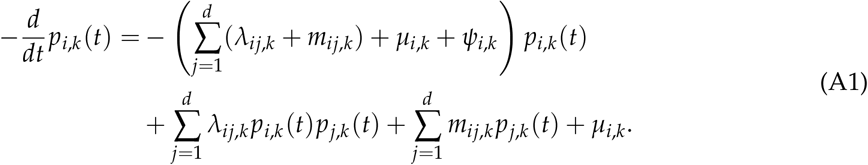

The terms on the right-hand side in the first line correspond to the probability of no event happening and the terms in the second line to either a branching, migration or death event happening. For a detailed derivation via Master equations, refer to Maddison *et al*. [14] or Stadler and Bonhoeffer [15].

For *t* = *t*_*k*_, *k* ∈ {0 … *n*}, we have,

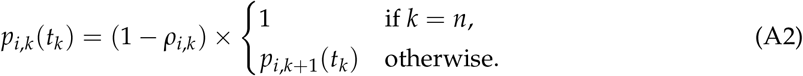

### Appendix A.2. Probability density of a sample-typed subtree

We use 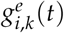 to denote the probability density that an individual represented by branch *e* at time *t* (with *t*_*k*−1_ ≤ *t* ≤ *t*_*k*_, *k* ∈ {1 … *n*}) in state *i* ∈ {1, …, *d*} has evolved between *t* and *T* as observed in the tree. Branch *e* is connected to two nodes, and we denote the more recent node with *n*_*e*_, occurring at time *t*_*k*−1_ < *t*_*e*_ ≤ *t*_*k*_. Node *n*_*e*_ represents either a sampling event (Figure A1a), a branching event (Figure A1b), or a degree-2 node at *t*_*e*_ = *t*_*k*_ with or without sampling (Figure A1c and d). One can show that 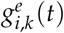 for *t* < *t*_*e*_ satisfies the ODE

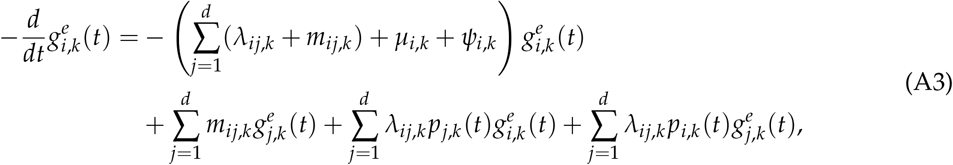

with the derivation being analogous to that of *p*_*i,k*_(*t*).

**Figure A1.**
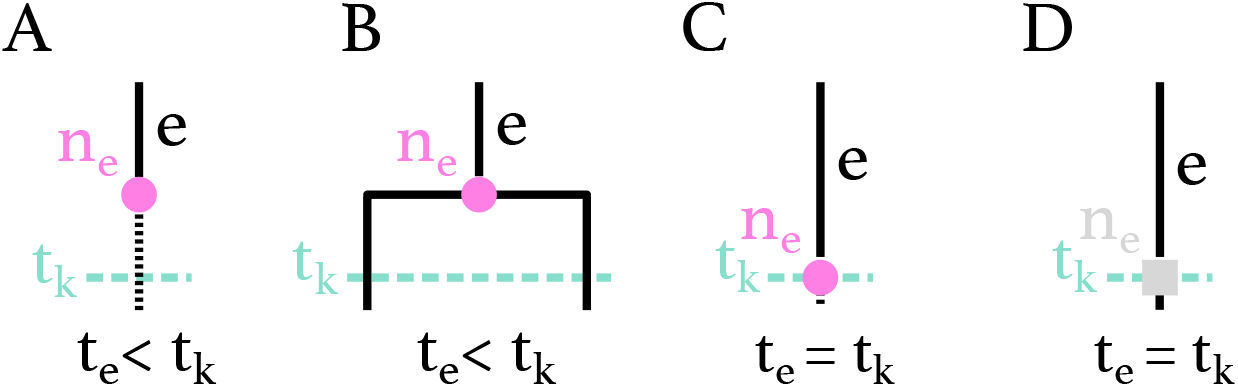
Possible configurations for node *n*_*e*_ on branch *e* at time *t*_*k*−1_ < *t*_*e*_ ≤ *t*_*k*_. (**a**) *n*_*e*_ is a *ψ*-sampled node at time *t*_*e*_ < *t*_*k*_, with or without sampled descendants. (**b**) *n*_*e*_ is a branching event at time *t*_*e*_ < *t*_*k*_. (**c**) *n*_*e*_ is a *ρ*-sampling node at time *t*_*e*_ = *t*_*k*_, with or without sampled descendants. (**d**) *n*_*e*_ is a degree-2 node at time *t*_*e*_ = *t*_*k*_ without sampling.

We denote the branch (resp. two branches) descending from *n*_*e*_ at time *t*_*e*_ with *e*_1_ (resp. *e*_1_ and *e*_2_). The initial conditions for the differential equations at node *n*_*e*_, i.e. the values of the probability densities at the most recent end of the branch *e*, are as follows:

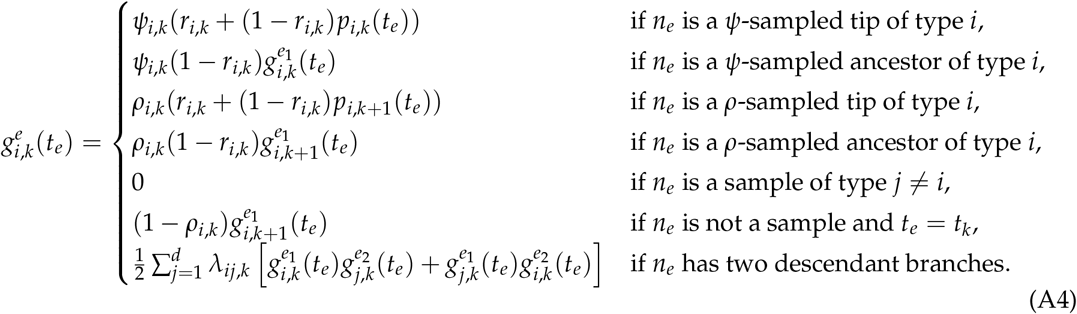

The 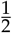 in the last equation is needed since we compute the probability density of an oriented tree.

### Appendix A.3. Probability density of a branch-typed subtree

When inferring branch-typed trees, we condition on the type of a branch at all times. Thus, we do not integrate over migration events or unobserved branching events changing the type of the tree lineage. We define *η*_*i,j*_ = 1 for *i* ≠ *j* and *η*_*i,j*_ = 2 for *i* = *j*. Equation A3 is replaced by,

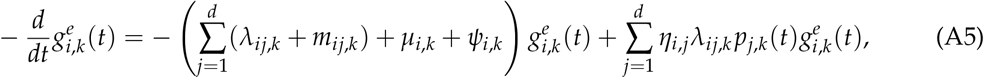

with initial conditions

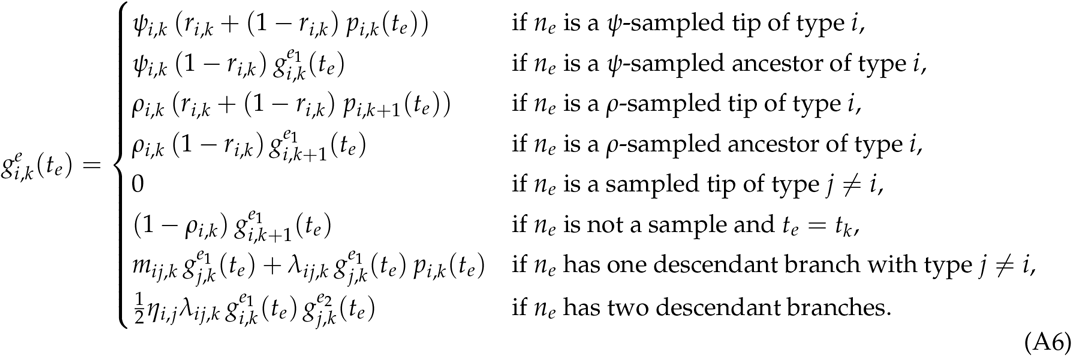

Here also, the 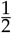 in the last equation is needed since we compute the probability density of an oriented tree. In branch-typed trees, a branch is always of one single type. Given the type of branch *e* is *I* implies that 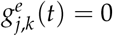 for *j* ≠ *i*. Indeed, Eqn. A6 states 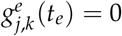 for *i* ≠ *j*. Further, Eqn. A5 specifies 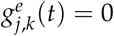 for all *t* < *t*_*e*_.

### Appendix A.4. Probability density of a sampled tree

The probability density of a sampled tree, with the lineage at time *t* = 0 being of type *i* and the branch being labelled with *e*, is the product of the probability density that the individual evolved as observed in the tree 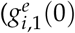 and the probability *h*_*i*_ that the individual at the start of the process is in type *i*.

Hence, the probability density of an oriented sampled tree 𝒯 under the multi-type birth–death model is

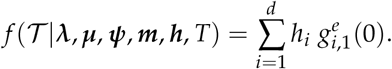

In BEAST 2, we infer labelled sampled trees, thus we need to calculate the probability of a labelled sampled tree. In a labelled sampled tree, each sample has a unique label, and orientations at branching events are ignored. In order to obtain the probability density of a labelled sampled tree, we need to transform the oriented tree probability density by multiplying by 2^*M*^ /*N*! where *M* is the number of branching events in the tree and *N* is the number of samples [20]. The probability density of a labelled tree 𝒯 under the multi-type birth–death model is thus

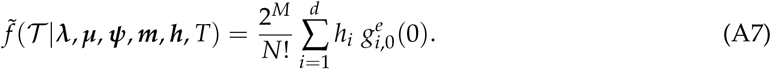

## Appendix B. Improved implementation of the tree probability density evaluation

The core component of the *bdmm* package is the evaluation of the tree probability density given by Eq. (A7). This involves numerical integration to solve the system of ODEs defined by Eqs. (A1)–(A2) and either Eqs. (A3)–(A4) (for sample-typed trees) or Eqs. (A5)–(A6) (for branch-typed trees).

In what follows, we discuss the numerical stability of our calculations.

### Appendix B.1. Extended numerical representation

The traditional floating point representation of a real number *x* is the closest number 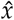 to *x*, where

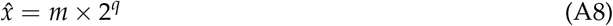

and where *m* (the *mantissa*) and *q* (the *exponent*) are signed binary integers of a specified number of bits. Note that the mantissa is understood to represent a fixed-precision number between 1 and 2. In the standard 64 bit double-precision floating point (DPFP) used by the original *bdmm* implementation [17], *m* is restricted to 53 bits and *q* is restricted to 11 bits. Ignoring the mantissa, this implies that the smallest representable non-zero absolute value is on the order of 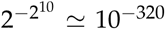. We calculate a probability density 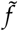 over tree space (together with intermediate values 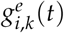 as part of the ODE integration), and these values can fall well below the lower limit imposed by the standard 64 bit DPFP representation, resulting in underflow errors.

To work around this limitation, our new implementation of *bdmm* employs a 96 bit extended precision floating point representation (EPFP) in which the mantissa is represented using a standard 64 bit double precision floating point number and the exponent is represented using a 32 bit signed integer value. This dramatically extends the range of possible values, in particular the smallest non-zero absolute value is 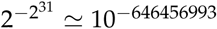.

A naive approach to employing the EPFP representation would be to use numbers of this kind at each and every step in the probability density calculation. This would ensure that all intermediate calculation results were accurately stored and would allow for accurate calculations at the next step. However, this approach has two main drawbacks. Firstly, existing numerical integration libraries almost exclusively use the DFPF representation. While integration algorithms could certainly be implemented for other representations such as the EPFP, this would be extremely time-consuming. Secondly, since DPFP calculations are implemented at a hardware level on modern processors, calculations using this representation are done very efficiently. Abandoning this primitive data type thus makes basic calculation steps considerably less efficient.

For these reasons, our approach involves a mixture using both representations. Numerical integration of the ODEs along each branch of the sampled phylogeny (Eqns. A3 and A5) is performed using the DPFP representation, while the combination of these integration results – done to provide the boundary condition for the next integration (Eqns. A4 and A6) – is achieved using the EPFP representation.

In order to avoid underflow at all times, the EPFP numbers are scaled before they are converted to the DPFP representation used by the integrator. The scaling procedure amounts to an integration by substitution. We scale 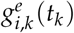 by the linear substitution function 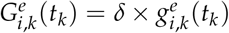, with *δ* being a scale factor. Since the original equations (A3) and (A5) are both linear in 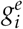, this scaling does not affect the results of their integration once the inverse scaling at time *t*_*k*−1_ is done. The method employed to choose an appropriate scale factor is described in detail below. The goal of scaling is to make sure all values stored as DPFP actually fit the window of values this representation can express. After each numerical integration step, results are converted back to EPFP and rescaled accordingly. The differential equation for the probability *p*_*i,k*_(*t*) of having no sampled descendants is not scaled at all as its integration does not cause any underflows in practice.

### Appendix B.2. Choice of a scale factor

The scale factor *δ* is shared among all subpopulations. Therefore, it has to be carefully chosen, so that all initial conditions fit in the window of values that can be represented in DPFP. To choose *δ*, we use three rules: A, B and C. We apply rule A if possible, otherwise rule B, and otherwise rule C. These rules are described below and illustrated in Figure A2, with a brief summary in legend.

Let 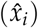 represent a set of floating point representations of real positive numbers (*x*_*i*_), as defined in Eq. A8. 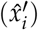 are the scaled values:

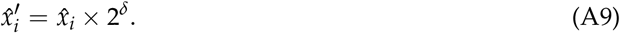

*q*_*min*_ and *q*_*max*_ denote the minimum and maximum exponents over all 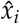. We define *q*_*low*_ = −1022 and *q*_*high*_ = 1023, respectively the lowest and highest acceptable exponents for a scaled value 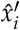. We define *q*_*gap*_ = 2040 the largest accepted difference between exponents across pairs of scaled values 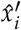. The values *q*_*low*_, *q*_*high*_ and *q*_*gap*_ were chosen to accommodate 64 bit DPFP representation, with some wiggle room to eliminate the need to account for edge cases.

**Rule A** is applied if:

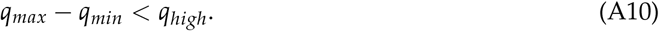

In this case, the scale factor *δ* is simply −*q*_*min*_. With this rule, the minimal exponent of scaled values becomes zero. Thus, scaled values are located between 1 and the maximum value in DPFP representation.

**Rule B** is applied if:

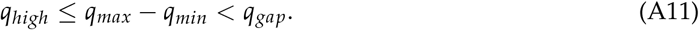

The scale factor is *δ* = *q*_*low*_ − *q*_*min*_ + (*q*_*gap*_ − (*q*_*max*_ − *q*_*min*_))/2. With this rule, exponents of scaled values are approximately centered around zero. Sets of values 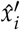 with greater variation between the smallest and biggest elements can be handled than with rule A.

In the rare case that neither rule A nor rule B can be applied, **rule C** is applied. All 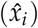 values with exponents *q*_*i*_ such that:

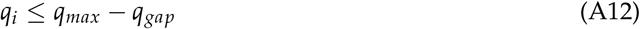

are set to zero. These are the smallest values among 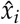 values. Then, rule B is applied to all non-zero 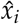 values.

### Appendix B.3. Performance improvements

We use various techniques to increase the efficiency of the numerical calculations performed by *bdmm*.

Prior to the integration of the coupled differential equations *p*_*i,k*_ and 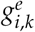 backwards in time, we calculate the values of *p*_*i,k*_ for every sampling event in the tree, by numerically integrating the ODEs for *p*_*i,k*_. We store the values obtained and use them when calculating the initial conditions for 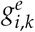 for every tip.

We also implement a parallelization scheme. An initial recursive tree traversal step is necessary to reach the tips of the tree, before launching numerical integration of the system of ODEs on 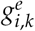 and *p*_*i,k*_ along the tree branches. During this traversal, when a node is reached whose left and right child subtrees are of significant size compared to the total tree size, a new computation thread is spawned and assigned to the traversal of one of the two subtrees. The initial thread continues onward with the traversal of the other one. This split between two threads is only executed when both subtrees represent more than a user-defined fraction of the total tree length, by default a tenth. This is done to prevent excessive numbers of threads from being created, since thread creation itself carries a computational overhead.

Finally, we replace the fixed-step size Runge-Kutta integrator used as default integrator in the original implementation by the fifth-order Dormand-Prince integrator for ODEs [27]. This integrator uses stepsize control, which improves the efficiency and accuracy of numerical integration steps. We use the existing implementation of these integrators from the *Apache Commons Math* Java library [28].

**Figure A2.**
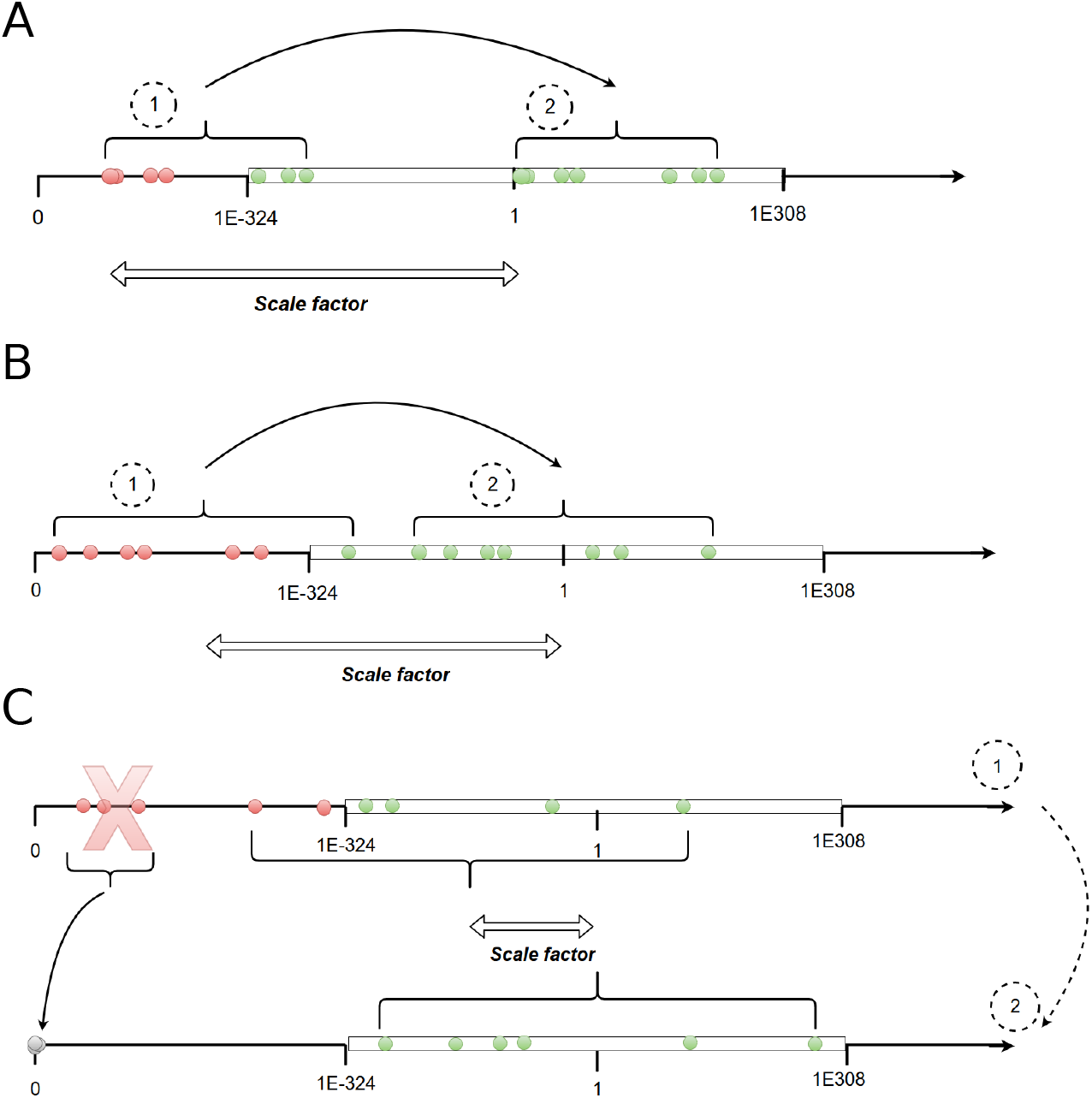
Scale factor choice. (**a**) Simplest case. The scale factor is the inverse of the smallest non-scaled value. (**b**) If A is not applicable, the scale factor is chosen such that the initial conditions are centered inside the range of values acceptable. The mid-point (on a log-scale) of this interval is approximately 1. (**c**) Last case, if all scaled values cannot fit at once inside the range of accepted values, the lowest non-scaled values are dropped and set to zero so that the problem simplifies to case 1 (panel **a**) or 2 (panel **b**). In all panels, the white rectangle represents values that can be represented using DPFP. Dots represents the values of initial conditions for the differential equations of the multi-type birth-death model, before (1) and after (2) scaling. Red dots represent values that are initially outside the window of values that can be represented using DPFP.

## Appendix C. Details on likelihood comparisons

To compare the results of computations performed with each *bdmm* version, we randomly-simulated sampled trees with 2 demes with fixed multi-type birth–death parameter values using MASTER [29]. We then computed tree likelihood values, varying one of the multi-type birth–deathparameters (either the birth or death rate of deme 1, corresponding to the tree origin location). Parameter values used for simulation and likelihood computation are listed in Table. A2. In Fig. 3 and Fig. A3, we show results for three ten-tip trees, for two of them we vary *λ*_1_ and for the third, we vary *μ*_1_. In Fig. A3, we also show results with a hundred-tip tree, varying *λ*_1_. For all simulated trees, likelihood results are identical between the original and new *bdmm* versions.

## Appendix D. Additional details on influenza data analysis

As we deal with pathogen sequence data, we adopt the epidemiological parametrization of the multi-type birth-death model as detailed in Kühnert *et al*. [17]. The epidemiological parametrization substitutes birth, death and sampling rates with effective reproduction numbers within types, rate at which hosts become noninfectious and sampling proportions.

For *i* ∈ {1, …, *d*} and k ∈ {1, …, n},

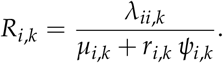

The rate of becoming uninfectious *δ*_*i,k*_ represents the inverse of the mean duration of infection:

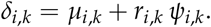

Based on our data, *ρ*_*i,k*_ = 0 for all *i, k*, as there is no singular point in time when a population-wide sampling effort was carried out which lead to multiple simultaneous samples. Thus, the probability for an individual to be sampled, or sampling proportion, is:

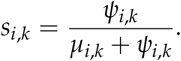

We assume *r* = 1 for all *i, k* in our analyses, i.e. individuals become non-infectious upon sampling. Further, we assume that *δ* is constant across locations *i* and time intervals *k*. Sampling proportion and migration are assumed not to change through time.

To study the seasonal dynamics of the global epidemic, we allow the effective reproduction number *R* to vary through time. To do so, we subdivide time into six-month intervals (starting April 1st and October 1st) for the time period during which we have samples. We set all values corresponding to the same season across different years to be equal for a particular location. Therefore, we infer two different values of *R* for each location *X*, one which corresponds to the April-September period: 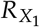 and another one for the October-March period: 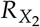. The location-specific sampling proportions *s*_*i*_ are assumed to be constant in the time interval in which we have samples, and null before the first sample.

Migration rates between types are inferred as products of a unique migration factor *σ*_*m*_ and relative rates *M*_*i,j*_:

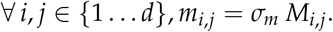

This setting allows us to use rather informative priors around 1 for the relative rates, and only requires a less informative prior for the unique migration factor. Table A3 lists the prior distributions assumed for the birth-death parameters.

Following [19], we use a GTR+Γ substitution model [30] and a strict molecular clock with the same priors as in [19].

The BEAST 2 analysis infers the birth-death model parameters, the substitution model parameters, and the clock model parameters together with the phylogenetic tree. In our analysis, the inferred phylogenetic trees are sample-typed trees. Thus, we do not attempt to reconstruct the history of migrations, rather we marginalize over all the possible migration histories. This is done to allow the MCMC chain to converge in shorter time compared to an analysis inferring branch-typed trees.

For each data analysis, we ran 10 parallel MCMC chains for 9 × 10^7^ steps each using the new implementation of *bdmm* as a package of BEAST 2.6 [16,31]. The computations were run on the Euler computation cluster at ETH Zürich. Each MCMC chain in the 175-sample analysis required 150 computation hours on average (1500 hours in total for ten chains). In the 500-sample analysis, each MCMC required 360 computation hours on average (3600 hours for ten chains). We used Tracer v1.6 [32] to check for convergence. The effective sample size was above 200 for all parameters. We combined the chains run in parallel into one using LogCombiner. We obtained the MCC tree using TreeAnnotator. Both LogCombiner and TreeAnnotator are available as part of BEAST 2.6. Finally, we plotted the numerical results with the R package *ggplot2* [33] and the MCC tree with *ggtree* [34].

**Table A1.** Distributions from which parameters were sampled for simulating trees. All parameters are constant through time.

| Parameter | Distribution |
| --- | --- |
| $\lambda_i$ | Unif(1, 3) |
| $\mu_i$ | $\lambda_i \times \text{Unif}(0, 1)$ |
| $m_{i,j}$ | Unif(0, 0.5) |
| $\psi_i$ | Unif(0.05, 0.5) |
| $r_i$ | Unif(0, 1) |

**Table A2.** Fixed parameter values for tree simulation and likelihood computations.

| Parameter | Value |
| --- | --- |
| $\lambda_1$ | 0.4 |
| $\lambda_2$ | 0.3 |
| $\mu_1$ | 0.27 |
| $\mu_2$ | 0.17 |
| $\psi_1$ | 0.03 |
| $\psi_2$ | 0.03 |
| $m_{1,2}$ | 0.03 |
| $m_{2,1}$ | 0.03 |
| Initial root state | 1 |

## Appendix E. Tables

## Appendix F. Additional figures

**Table A3.** Prior distributions for parameters of the multi-type birth-death model in the seasonal influenza analysis.

| Parameter | Prior distribution |
| --- | --- |
| $R$ | LogNormal(0, 1.0) |
| $\delta$ | LogNormal(4.5, 0.15) |
| $\sigma_m$ | LogNormal(0, 2.0) |
| $M_{i,j}$ | LogNormal(0, 0.5) |
| $s$ | Exp(0.001) truncated on $[0, 1]$ |
| $r$ | Beta(10.0, 1.5) |
| $T$ | LogNormal(2.0, 1.0) |

**Figure A3.**
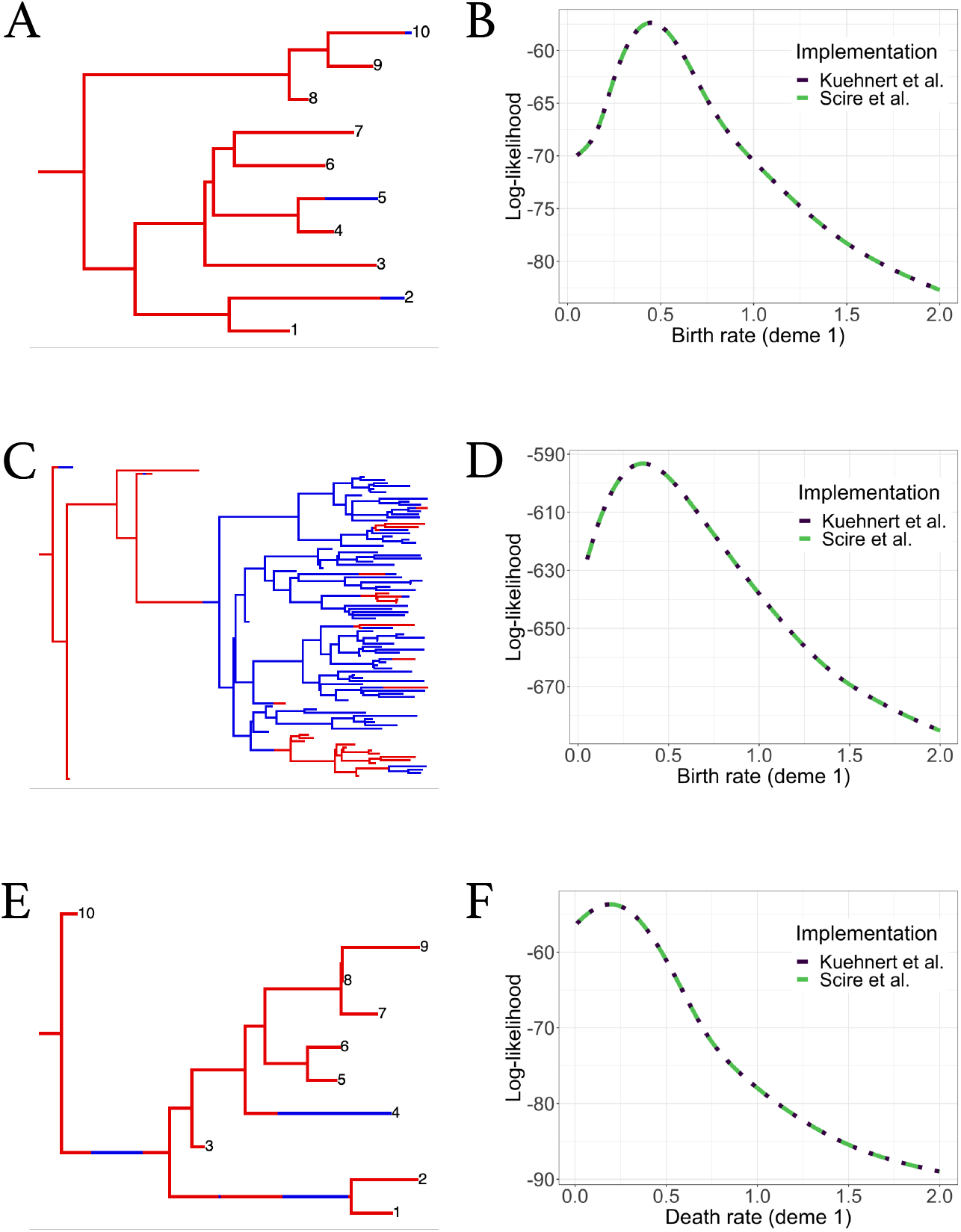
Comparisons of likelihood computation results between the original and improved *bdmm* versions for additional trees. (**a-b**) Randomly-simulated ten-tip tree and log-likelihood computation results against *λ*_1_ (birth rate of red deme). (**c-d**) Randomly simulated hundred-tip tree and log-likelihood computation results against *λ*_1_ (birth rate of red deme). (**e-f**) Randomly-suimulated ten-tip tree and log-likelihood computation results against *μ*_1_ (death rate of red deme).

**Table A4.** Inferred parameter values for Influenza A virus analysis under the multi-type birth-death model. For each parameter, the lower and upper bounds for the 95% Highest Posterior Density interval (hpd_low, hpd_high) are given along with the median (m). *N, T*, and *S* refer respectively to the North, South and Tropics. For effective reproduction numbers *R*, the first subscript is the location while the second one refers to the period of the year. *s, w, as, om* respectively refer to *summer, winter, april-september*, and *october-march*. Thus, for instance, 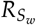 refers to the effective reproduction number of samples from the southern hemisphere during the winter season. The tree height *t* is given in number of years, *M* is given in migrations/lineage/year and *δ* is given in years^−1^. The remaining parameters are unitless.

|  | 175 samples |  |  | 500 samples |  |  |
| --- | --- | --- | --- | --- | --- | --- |
|  | m | hpd_low | hpd_high | m | hpd_low | hpd_high |
| $t$ | 3.342 | 3.048 | 3.643 | 6.645 | 6.313 | 7.031 |
| $\delta$ | 90.464 | 75.836 | 105.207 | 101.582 | 87.197 | 116.768 |
| $R_{N_s}$ | 0.354 | 0.199 | 0.556 | 0.505 | 0.263 | 0.815 |
| $R_{N_w}$ | 0.971 | 0.927 | 1.01 | 1.001 | 0.98 | 1.02 |
| $R_{T_{as}}$ | 1.048 | 1.026 | 1.071 | 1.005 | 0.992 | 1.017 |
| $R_{T_{om}}$ | 0.991 | 0.969 | 1.013 | 1.01 | 0.998 | 1.022 |
| $R_{S_s}$ | 0.558 | 0.335 | 0.783 | 0.774 | 0.679 | 0.861 |
| $R_{S_w}$ | 1.08 | 1.037 | 1.123 | 1.027 | 1.002 | 1.051 |
| $\sigma_m$ | 0.475 | 0.196 | 0.869 | 0.304 | 0.147 | 0.524 |
| $M_{N,T}$ | 0.871 | 0.245 | 1.923 | 1.064 | 0.3 | 2.422 |
| $M_{N,S}$ | 0.894 | 0.253 | 1.965 | 0.838 | 0.264 | 1.744 |
| $M_{T,N}$ | 2.183 | 0.877 | 4.174 | 1.568 | 0.635 | 2.973 |
| $M_{T,S}$ | 0.561 | 0.205 | 1.08 | 0.694 | 0.332 | 1.248 |
| $M_{S,N}$ | 1.001 | 0.273 | 2.261 | 0.887 | 0.275 | 1.93 |
| $M_{S,T}$ | 1.005 | 0.279 | 2.173 | 1.13 | 0.368 | 2.468 |
| $h_N$ | 0.292 | 0 | 0.777 | 0.296 | 0 | 0.779 |
| $h_T$ | 0.3 | 0 | 0.784 | 0.292 | 0 | 0.777 |
| $h_S$ | 0.283 | 0 | 0.769 | 0.289 | 0 | 0.767 |
| $P(\text{root in N})$ | 0.024 | 0 | 0.535 | 0.023 | 0 | 0.501 |
| $P(\text{root in T})$ | 0.849 | 0.167 | 1 | 0.863 | 0.176 | 1 |
| $P(\text{root in S})$ | 0.041 | 0 | 0.731 | 0.044 | 0 | 0.67 |

